# Chronic hypoxia regulates Cytoplasmic Polyadenylation Element Binding Protein 2 alternative splicing to promote HIF1a translation

**DOI:** 10.1101/2020.10.05.325290

**Authors:** Emily M. Mayo, Shaun C. Stevens, Anika N. Ali, Christina J. Moss, Sean P. Lund, Gina S. Nazario-Muñoz, Jennifer B. Permuth, Sandy D. Westerheide, Charles E. Chalfant, Margaret A. Park

## Abstract

HIF1 (Hypoxia-inducible Factor 1) is a transcription factor that plays a crucial role in the hypoxia stress response. However, chronic hypoxia exposure can cause irreversible physiological changes that can lead to pulmonary hypertension (PH), and the need for therapeutics to ameliorate these conditions is great and unmet. Previous studies in our lab have demonstrated that CPEB2 (cytoplasmic polyadenylation element binding protein 2) is a translational repressor of one of the HIF1 subunits: HIF1α. Our lab demonstrated that the alternatively spliced CPEB2A isoform of CPEB2 is a repressor of translation, while the CPEB2B isoform is a translational activator of HIF1α during hypoxia, suggesting a major regulatory role for CPEB2 AS in the pulmonary hypoxic response. Although it is well established that during hypoxia, HIF1α levels are dramatically upregulated due to a decrease in the degradation of this factor, we propose that during chronic hypoxia, the expression of HIF1α is maintained via a translational mechanism, likely alongside a decrease in proteolytic degradation. In this study we demonstrate that depletion of the CPEB2B splice isoform has an inhibitory effect on the translation of nascent HIF1α protein during chronic hypoxia, but not the acute phase. We further demonstrate this pathway is dependent on the initiation factor eIF3H. Finally, we show data that indicate CPEB2A and CPEB2B bind differentially to cytoplasmic polyadenylation element consensus sequences depending on the surrounding sequence context. These findings are important since they provide evidence for the potential of CPEB2 to act as a therapeutic target for treating chronic hypoxia-related pulmonary diseases.

## INTRODUCTION

When hypoxic conditions are encountered, the human pulmonary system undergoes alterations in gene and protein expression in order to adapt to an oxygen deprived environment. Under the control of hypoxia-inducible factors (HIF proteins), the transient expression of angiogenic and vasoconstrictive genes can help attenuate the hypoxic stress response. However, over longer exposures, expression of HIF-dependent genes can lead to detrimental and irreversible effects such as rearrangement of the lung vasculature and eventually occlusion of blood vessels (**1**,**2)** As an example, one of the most common causes of chronic hypoxia (CH) in the lungs is chronic obstructive pulmonary disease (COPD), commonly induced by smoking and accounting for as many as 8 out of 10 COPD-related deaths **(3)**. As a more timely example, COVID-19 patients often demonstrate hypoxia secondary to their infection in the lungs (**4**). In addition to COPD and COVID-19-mediated hypoxia, the chronic hypoxic response can be initiated by various pulmonary cancers, acute lung injury, sleep apnea, obesity and smoking. Treatment for these conditions has been limited to vasodilators and oral medications which, alongside hospital stays that cost around $110 billion US dollars annually to treat **(3**,**5)**.

To compensate for the poor oxygen flow throughout the lungs of a patient with chronic hypoxia, endothelial and smooth muscle cells throughout the pulmonary system proliferate, blood vessels constrict, and pulmonary blood pressure rises. These pathological, yet at first necessary changes ultimately lead to pulmonary hypertension and in the most severe cases, right-ventricular heart failure and death (**6**,**7**,**8**). Targeting symptoms at the source could potentially alleviate this cost and the morbidity associated with CH and it is thus important to understand how the CH response mechanism is regulated. To address gaps in current knowledge, our study aims to identify novel regulators of the hypoxic response with a focus on the chronic phase of the response, to open the door to new therapeutic targets.

Under normoxic conditions, the constitutively expressed HIF1β subunit remains in the cytosol, whereas a shift into hypoxia is required to increase expression of the short-lived HIF1α subunit **(9)**. The current reigning dogma for HIF1α expression is that during hypoxia, HIF1α levels increase due to a decrease in HIF1α ubiquitination. When oxygen is present, HIF1α is hydroxylated on conserved proline residues **(9**,**10)** causing its recognition by the von Hippel Lindau (VHL) ligase **(5**,**9)**. VHL tags HIF1α with ubiquitin, targeting it to the proteasome for degradation. However, as oxygen levels drop, HIF1α accumulates and binds to its partner, HIF1β, forming the HIF1 complex. This complex acts as a transcription factor to downstream targets **(11-13)**.

One such target of HIF1 is Vascular Endothelial Growth Factor (VEGF). Transcription of VEGF is regulated by the presence of a hypoxia response element (HRE) within the promoter of the VEGF transcript, which allows HIF-1 to both bind to the VEGF promoter and induces its expression during low oxygen conditions. Once transcribed, the binding of VEGF to its receptor (VEGFR) induces angiogenesis and promotes vasculogenesis (**14, 27**,**30-35**).

Although the ubiquitination/proteasomal degradation mechanism of HIF1α regulation is well established, our lab has observed prolonged HIF1α expression during the chronic phase of hypoxic stress, indicating a potential secondary mechanism that maintains HIF1α expression levels during chronic hypoxia. Published studies in our lab have demonstrated that the alternative splicing (A/S) of cytoplasmic polyadenylation element binding protein 2 (CPEB2) may play a role in HIF1α expression during the chronic hypoxic response (**15**). CPEBs are a family of proteins (CPEB1, CPEB2, CPEB3, and CPEB4) that generally act to suppress mRNA translation by modulating polyadenylation (**15**,**16**). Based on published work from our lab, the A and B isoforms of CPEB2 differ by a single 90 base-pair exon (**15**). This exon is excluded in the CPEB2A isoform, while it is included in the CPEB2B isoform. A splicing factor known as SRSF3 (arginine/serine-rich splicing factor 3) (**15, 17, 31**) binds to a consensus sequence near the 3’ end of the exon, forcing its inclusion in the final transcript of CPEB2B (**15, 17**). In this manuscript, we provide evidence that the B isoform of CPEB2 is required for the translation of new HIF1α protein specifically during the chronic stages of hypoxia exposure *in vitro*. Furthermore, we provide data indicating that the eIF3H subunit of the pre-initiation complex may aid CPEB2B in initiating HIF1α translation via preferential association with this protein.

Therefore, the alternative splicing of CPEB2 is a novel regulator of HIF1α during hypoxia. Future studies may be possible to target the expression of CPEB2B in COPD and other hypoxia-linked animal models and eventually aim to regulate HIF1 in patients suffering from chronic hypoxic disorders.

## RESULTS

### CPEB2 alternative splicing is dysregulated during chronic hypoxia

Seminal studies in the field have demonstrated that CPEB2 localizes via a low complexity sequence to stress granules which also contain mRNA transcripts (**32**). Generally, binding occurs at consensus sequences called cytoplasmic polyadenylation elements (CPE sites) **(18**,**19)**. These findings indicate that members of the CPEB family play a large role in stress responses, as the mRNA species found in stress granules are linked to stress processes (**15-19**). In light of the current literature and our previous findings, we assessed whether exposure to hypoxia would alter the CPEB2 A/B expression levels. For the purposes of these studies in a cell model, we define the acute hypoxic response as occurring prior to 24 hours and chronic exposure from 24-96 hours (the longest we have treated)(**20**). In general agreement with others’ findings, we found that HIF1α protein levels increased as early as 4 hours after exposure to hypoxic conditions *in vitro* (2% O_2_) and remained elevated well into 96 hours of constant exposure to hypoxic conditions. Correlative to these findings, the ratio of CPEB2A to CPEB2B dramatically decreased during later time points (**Fig. 1A**). Hence, we observed a reliable decrease in the CPEB2 A/B ratio as hypoxia progresses into the chronic stage (**Fig. 1A, S1**). Primary pulmonary arterial endothelial cells (hPAECs) demonstrated a similar pattern in both HIF1α protein and the CPEB2A/B ratio over the course of 96 hours (**Fig. 1B,F,G**). A Pearson’s correlation analysis demonstrates a negative correlation between CPEB2A/B ratio and HIF1α levels (correlation coefficient of -0.72; **Fig. 1C-E**).

**Figure 1.**
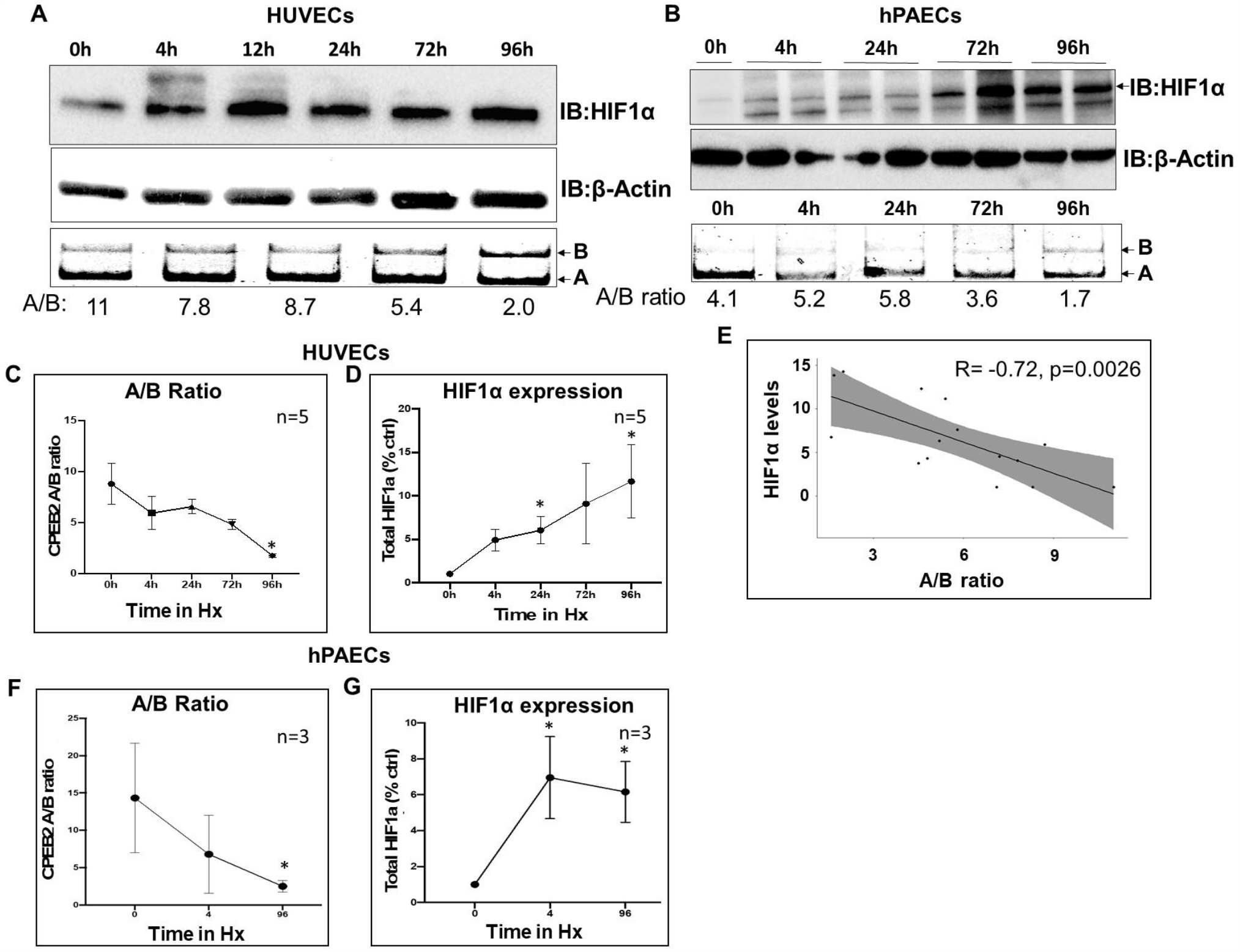
CPEB2 A/B RNA and HIF1a protein expression during hypoxia. (A) HUVECs (N=3: P3,P4,P6) and (B) PAEC’s (N=2: P3) were grown to 75-85% confluence before being placed in a hypoxia chamber (2% O_2_) for the indicated amount of time. The indicated antibodies were used after immunoblotting (top blots) or competitive RT-PCR was performed for CPEB2 AS (bottom gels). HUVEC results were densitometrically assessed for the average CPEB2 RNA A/B ratio (C) and HIF1α levels (D) and graphed for XY linear regression. (E) The A/B ratio/HIF1α levels were then correlated (Pearson’s R_2_ and p-value are indicated).

### CPEB2B regulates HIF1α protein levels during chronic hypoxia

As previously demonstrated by our lab ectopic expression of CPEB2A leads to a decrease in HIF1α protein expression, whereas ectopic expression of CPEB2B will lead to an opposing increase of this factor during normoxic conditions (**15**). Expanding on these findings, we assessed the impact of CPEB2B depletion on the expression of HIF1α during both acute and chronic hypoxia phases.

Our lab has established that the alternative splicing of CPEB2A is at least partially regulated by the *trans*-factor SRSF3 (Serine/Arginine-rich splicing factor 3) (**15-17**). Therefore, we used antisense oligonucleotides (ASOs) to block the binding site of SRSF3 to its consensus sequence in exon 4 of CPEB2B (**Fig.2A**). If SRSF3 cannot bind to its consensus sequence, exon 4 will be excluded from the final transcript and the ratio of CPEB2A/B will increase as shown in **Fig. 2C, 2F**. CPEB2B knockdown mediated by both antisense oligonucleotides (**Fig. 2C**) and siRNA **(Fig. 2B)** targeted towards CPEB2B (included exon) confirmed that HIF1α expression is in part dependent on CPEB2B expression (**Fig. 2B-C**) during the chronic phase of hypoxia. More specifically, we find that inhibition of CPEB2B at the later 72-96 h time point prevented HIF1α expression leading to expression values similar to those observed under normoxic conditions (**Fig. 2B-E**). Finally, we demonstrate in **Fig. 2H-M** that induction of endothelial sprouting (both cumulative sprout length (CSL) and total sprout number) in HUVEC organoids is inhibited by the prevention of CPEB2 exon inclusion. Interestingly, further findings indicate that ectopic expression of CPEB2B leads to increases in CSL but no significant differences in sprout number when compared to controls (**Fig. 2K-M**).

**Figure 2.**
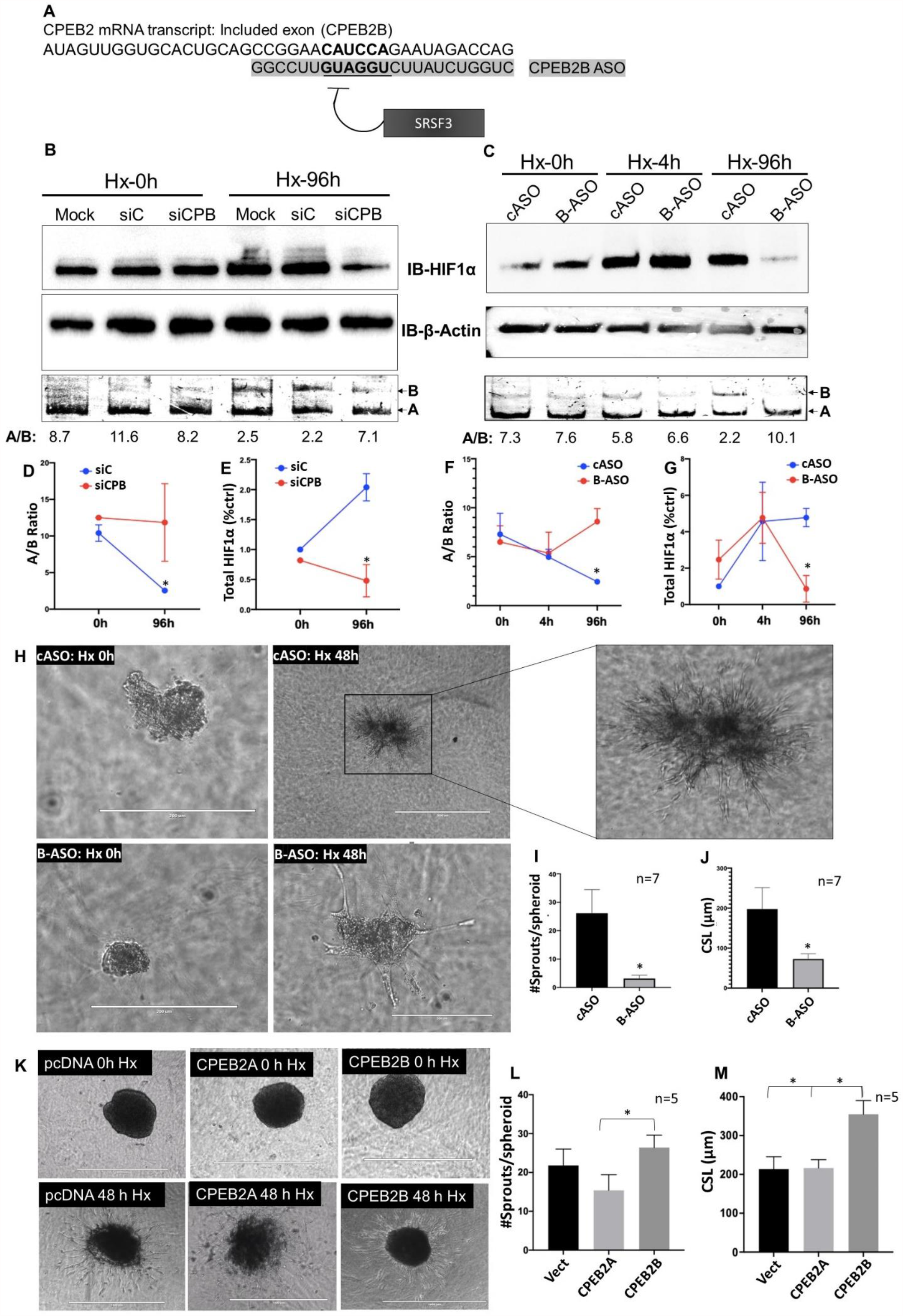
CPEB2B regulates HIF1α expression during Chronic Hypoxia. A) Schematic of the CPEB2B ASO design. (B-C) HUVECs (N=3: P3, P4) were subjected to hypoxia (2% O_2_) for the indicated times -/+ either mock conditions (Mock), control siRNA (siC) or siRNA targeted towards CPEB2B (siCPB), (B); or control ASO (c-ASO) or CPEB2B ASO (B-ASO) (pre-treated, C). Both RNA and protein were harvested and subjected to either immunoblot using the indicated antibodies (top blots) or was subjected to quantitative “splicing” RT-PCR for CPEB2 A/S (bottom gels). (D-G) Change in the CPEB2 A/B ratio (D,F) and HIF1α protein expression (E,G) were assessed via densitometry for depletion of CPEB2B either via siRNA (D,E) or ASO (F,G). (H-M) HUVEC spheroids were assessed via microscopy (H, K) and both number of sprouts per spheroid (I, L) and cumulative sprout length (CSL, J, M) were measured after either CPEB2B ASO depletion or in cells ectopically expressing either CPEB2A or CPEB2B (or vector control – Vect). The scale bar for ASO micrographs is 200 microns except for the control ASO spheroid at 48 h (the spheroids grow larger during sprouting, inset is a blown-up image to show sprout number). Scale bars for ectopic expression are 1000 microns. *indicates p < 0.05 via repeated measures ANOVA or t-test.

### HIF1α is synthesized at the translational level during Chronic Hypoxia

A large body of published data demonstrates that increases in HIF1α during hypoxia are due to a decrease in ubiquitination of this protein. However, published work in our lab has demonstrated that CPEB2 A/S plays a regulatory role in the *translation* of HIF1α (**15**). Hence, it is plausible that CPEB2 A/S allows the maintenance of enhanced HIF1α expression levels during later hours of hypoxia via nascent protein synthesis. Indeed, we find that inhibiting protein synthesis via cycloheximide induced a shift in HIF1α expression both in HUVEC’s and in hPAEC’s. After 4 hours of exposure, inhibition of translation had no significant effect on HIF1α expression, compared to 96 hours of exposure, where HIF1α protein levels were comparable to those observed during normoxia (**Fig. 3A,B; S2**). VEGF expression is also decreased at this time point after cycloheximide, both in HUVECs and hPAECs (data not shown). Interestingly, we find that cycloheximide induces HIF1α expression under normoxic conditions (**Fig.3A, S2**), perhaps due to the induction of translational stress, and that our data at 4 hours of hypoxia are somewhat variable. Hence, we assessed nascent protein production during short and long-term hypoxia exposure to confirm our findings.

**Figure 3.**
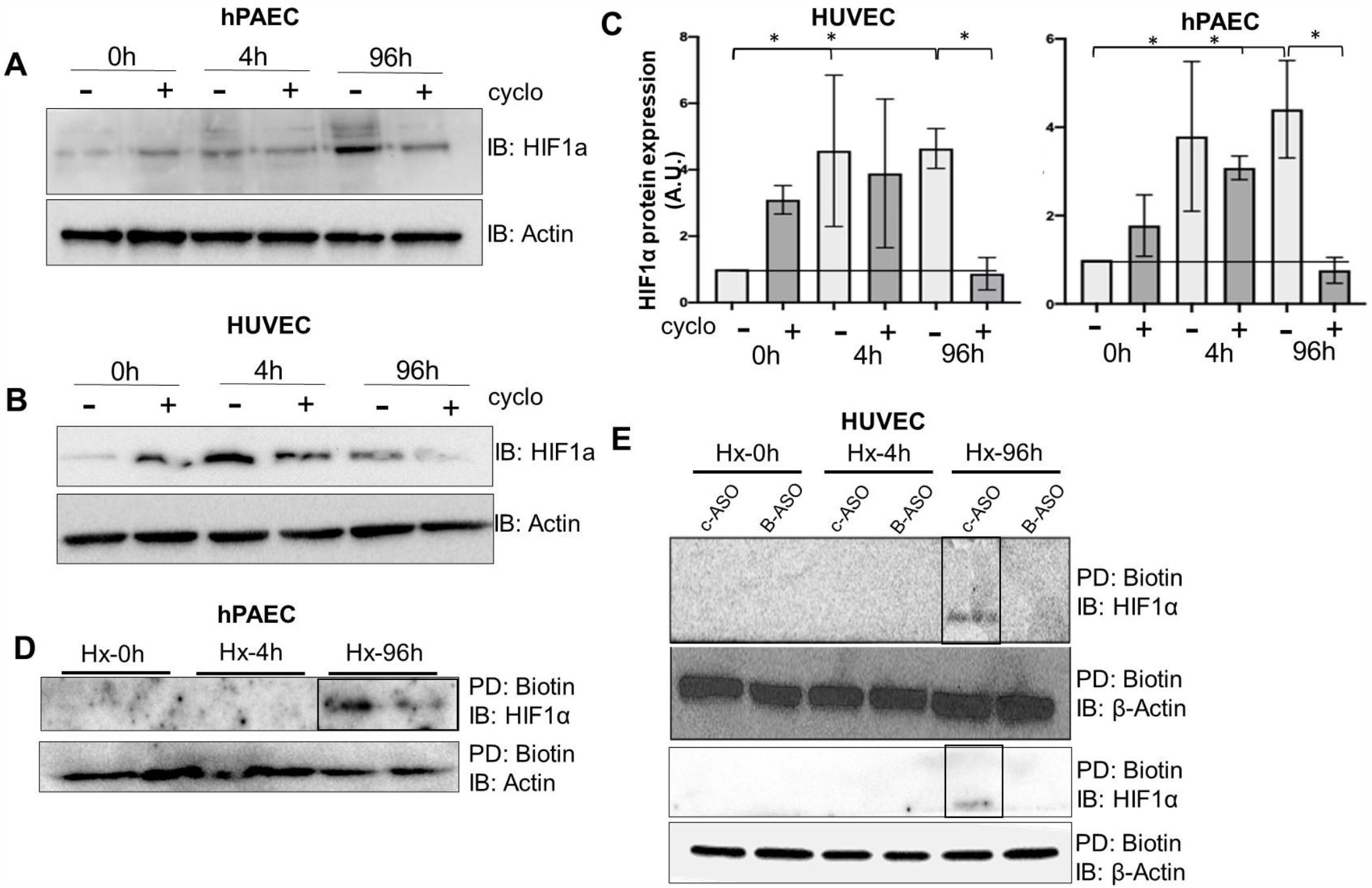
HIF1α levels are maintained by translation during chronic hypoxia. (A-C) HUVECS (P3-P6) and hPAECs (P=5) were incubated with cycloheximide (5ug/ml) 4 hours prior to harvesting and blotted with the indicated antibodies. HIF1α levels were assessed via ImageJ and graphed for total protein. (D-E) Nascent proteins were labelled with biotin, then precipitated using streptavidin (PD:Biotin). Labelled proteins were immunoblotted (IB) using the indicated antibodies. Blots in (E) and replicates in (D) represent biological replicates. Blots are representative of an n ≥3; *=p<0.05 via repeated measures ANOVA.

### CPEB2 alternative splicing regulates HIF1α translation during chronic hypoxia

In general agreement with our data in **Fig. 3A,B**, we used “CLICK” chemistry to detect nascent HIF1α protein via incubation with the methionine mimic with L-azido-homoalanine (AHA) and subsequent labeling with biotin (**Fig. 3C**). Nascent HIF1α protein was only observed at the 96 hour time point, but not the acute 4 hour time point. Nascent HIF1α expression was inhibited by the CPEB2B ASO, indicating that this pathway is dependent upon CPEB2B expression (**Fig. 3D-E**).

### CPEB2B induces downstream VEGF expression

When hypoxic conditions are encountered, the HIF1 transcription factor binds to hypoxic response elements within the promoter of transcriptional targets such as VEGF **(Fig. 4A)**. VEGF then induces vascular formation at the site of hypoxia. Therefore, we wished to establish whether targeting CPEB2 A/S can regulate downstream HIF1α targets linked to angiogenesis such as VEGF and thus we inhibited CPEB2B expression using B-ASO. We found that knocking down CPEB2B attenuates the expression of downstream targets of HIF1α, such as VEGF. Interestingly, forced expression of HIF1α “rescues” this effect indicating that HIF1α may lie downstream of CPEB2B in the pathway **(Fig. 4B-C)**.

**Figure 4.**
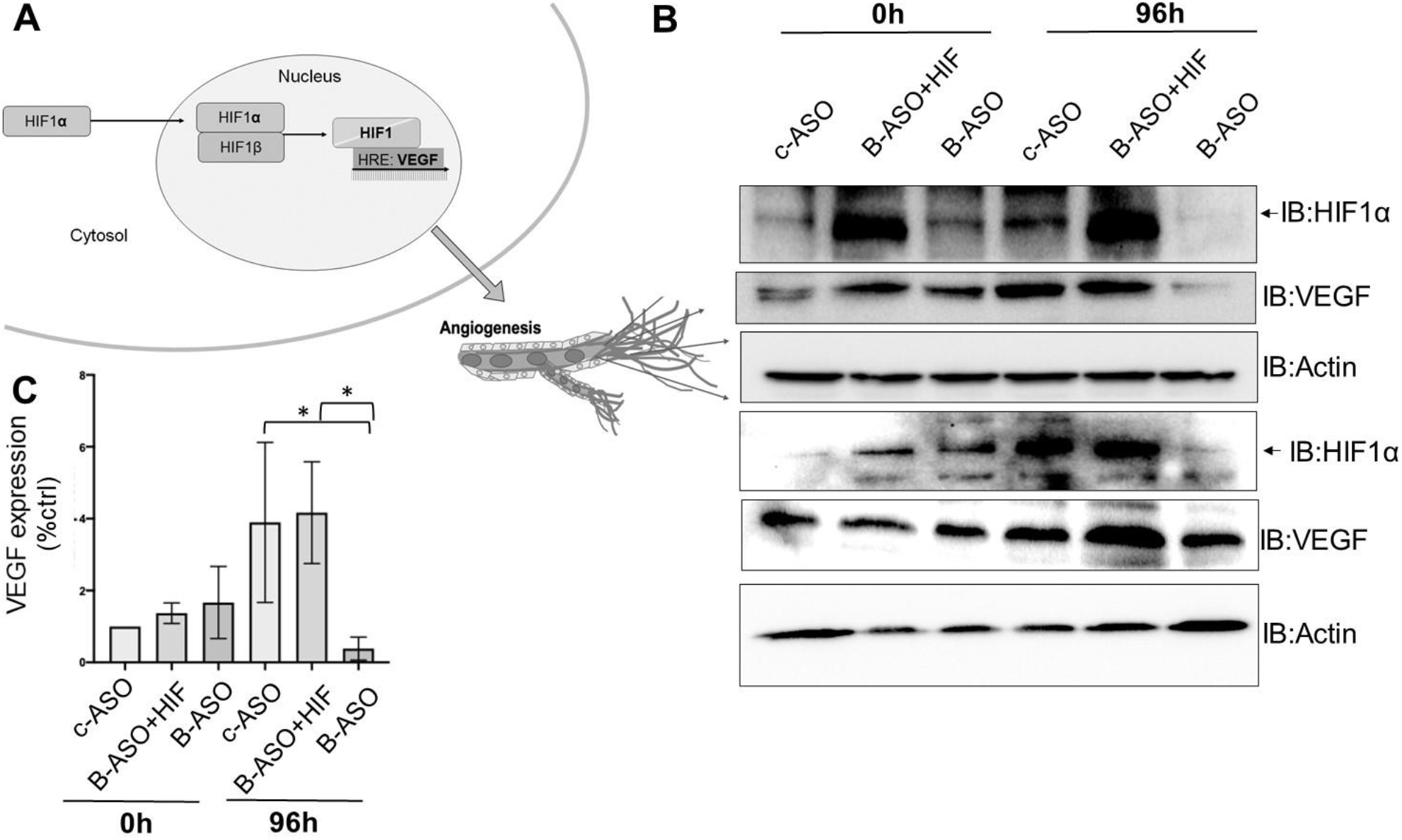
CPEB2B regulates downstream VEGF expression during chronic hypoxia. (A) Representation of HIF1α activity. (B-C) HUVECs were transfected with the indicated ASOs (control or CPEB2B) or a plasmid to express HIF1α, then exposed to hypoxia. Immunoblots were then performed with the indicated antibodies (repeat blots represent biological replicates). Densitometry was determined (shown in C). n≥3 for all assays; *=p<0.05 via ANOVA

### CPEB2B binds differentially to mRNA sequences and regulates HIF1α translation at least partially via eIF3H

To ascertain why CPEB2B function varies from the CPEB2A isoform, we assessed the RNA binding capabilities of the CPEB2 isoforms. Our results indicate that CPEB2A binds to RNA sequences in the HIF1α 3’UTR directly as demonstrated by a gel shift assay (**Fig.5A-B, S4**). We then wished to determine if CPEB2B binds differentially to this same site. We therefore used biotinylated RNA as “bait” to pull down immunoprecipitated CPEB2A or CPEB2B. Our findings indicate that CPEB2A binds robustly to sequences surrounding the CPE sites in the 3’UTR of both HIF1α and TWIST1, another RNA species with CPE sites in its 3’UTR. CPEB2B on the other hand, does not bind to the HIF1α CPE site as readily as CPEB2A (**Fig.5A,C, S4**). Further analysis of the HIF1α mRNA sequence indicates that there are other CPE consensus sequences in the coding region, specifically in exon 10 of the mRNA sequence. We used biotinylated sequences surrounding this sequence in binding assays using immunoprecipitated CPEB2A and B (see **Table 1**). We find that, while the A isoform binds more readily to the 3’UTR sequence, the B isoform binds preferentially to the sequence in exon 10 of HIF1α, indicating that inclusion of exon 4 in the CPEB2 protein may shift sequence specificity of this factor (**Fig. 5D**).

**Table 1.**
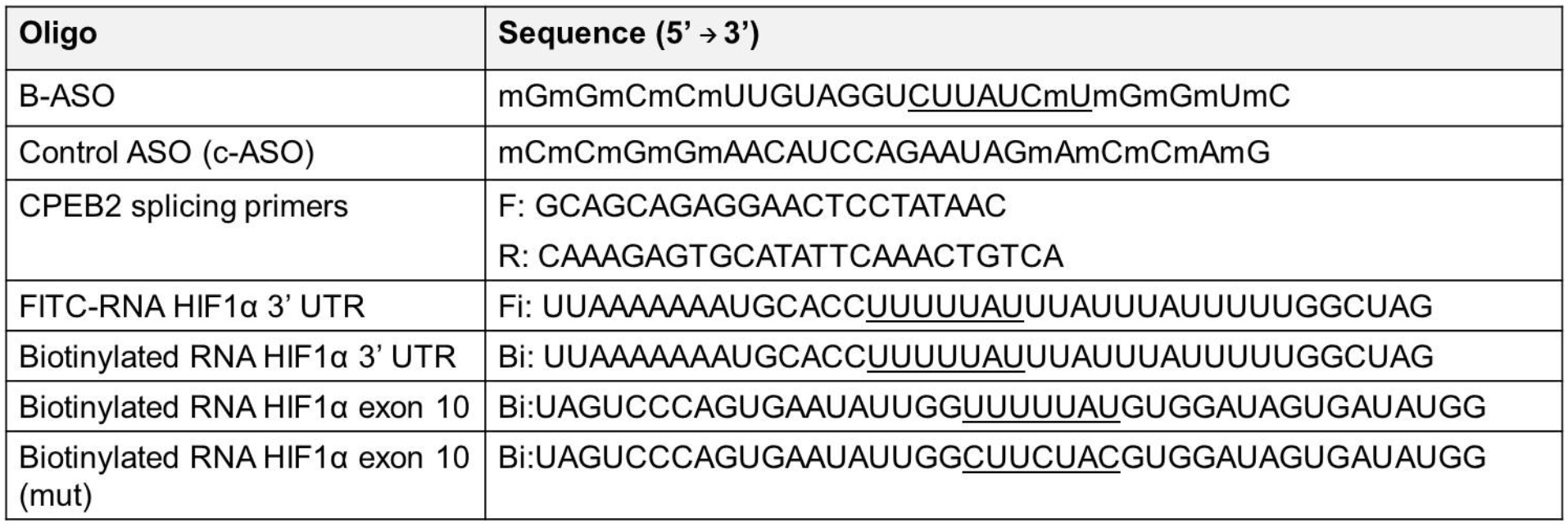
RNA and DNA Sequences.

**Figure 5.**
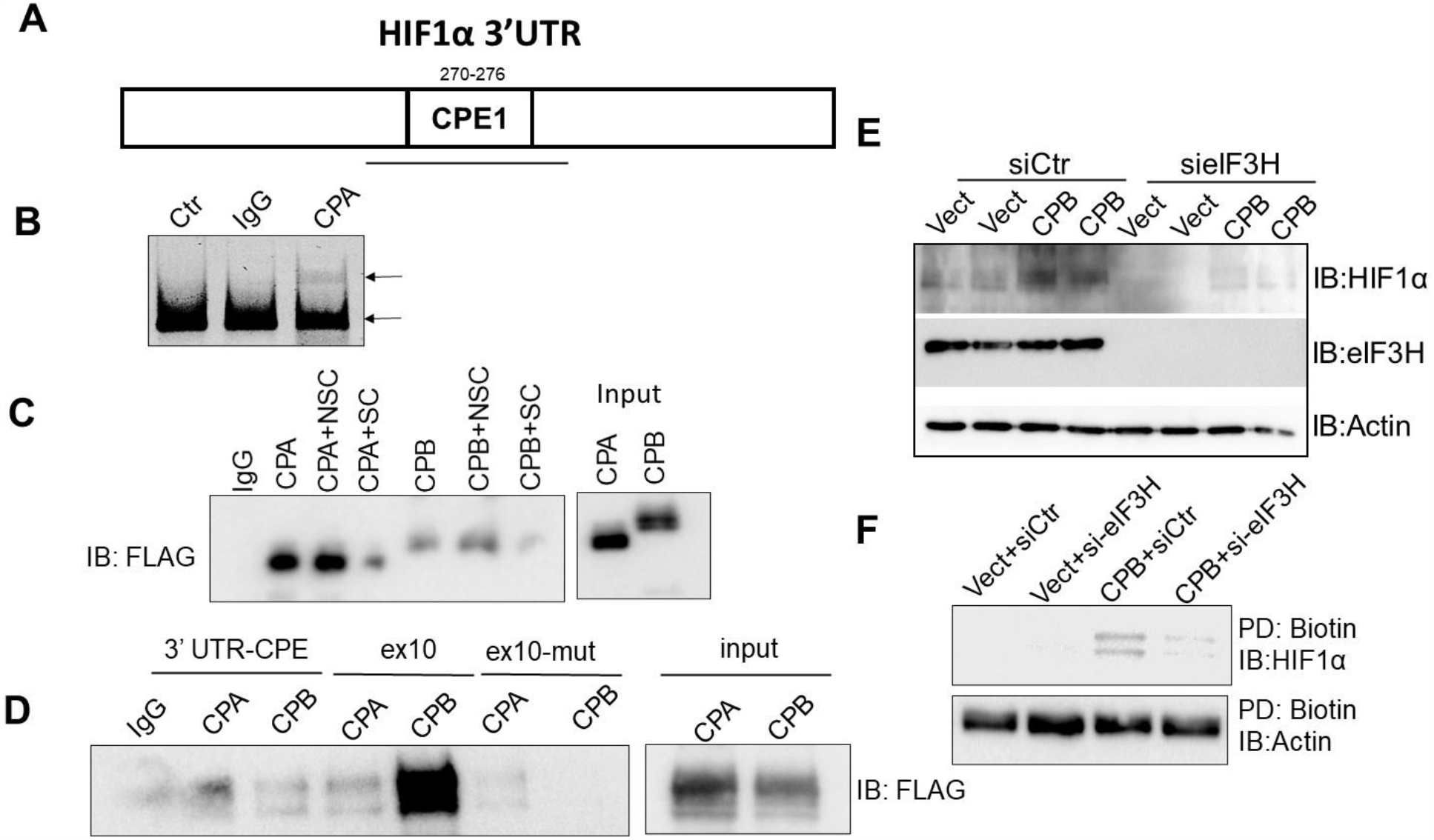
CPEB2B binds differentially to the 3’UTR region and to a region of HIF1α exon 10 and regulates HIF1α expression via eIF3H. (A) Schematic representation of the HIF1α 3’ UTR (CPE=cytoplasmic polyadenylation element, pA=alternative polyadenylation site). (B) FLAG-tagged CPEB2A was immunoprecipitated and exposed to FITC-labeled RNA sequences surrounding the HIF1α 3’ UTR CPE sites (CPE). Electrophoretic shift assay was performed. (C) Streptavidin-biotin affinity pull-down was performed using immuno-purified CPEB2A or CPEB2B with sequences corresponding to the 3’ UTR of HIF1α or TWIST1 published CPE sites (B) NSC=non-specific competitor, SC=specific competitor. (D) Streptavidin biotin affinity pulldown using either the 3’UTR CPE site, or sequences surrounding the CPE site in HIF1α exon 10. (E-F) HUVECs were transfected with published siRNA targeting eIF3H. Some samples were subjected to nascent protein labeling (F), then immunoblotted using the indicated antibodies (E-F). n≥3 for all assays.

**Figure 6:**
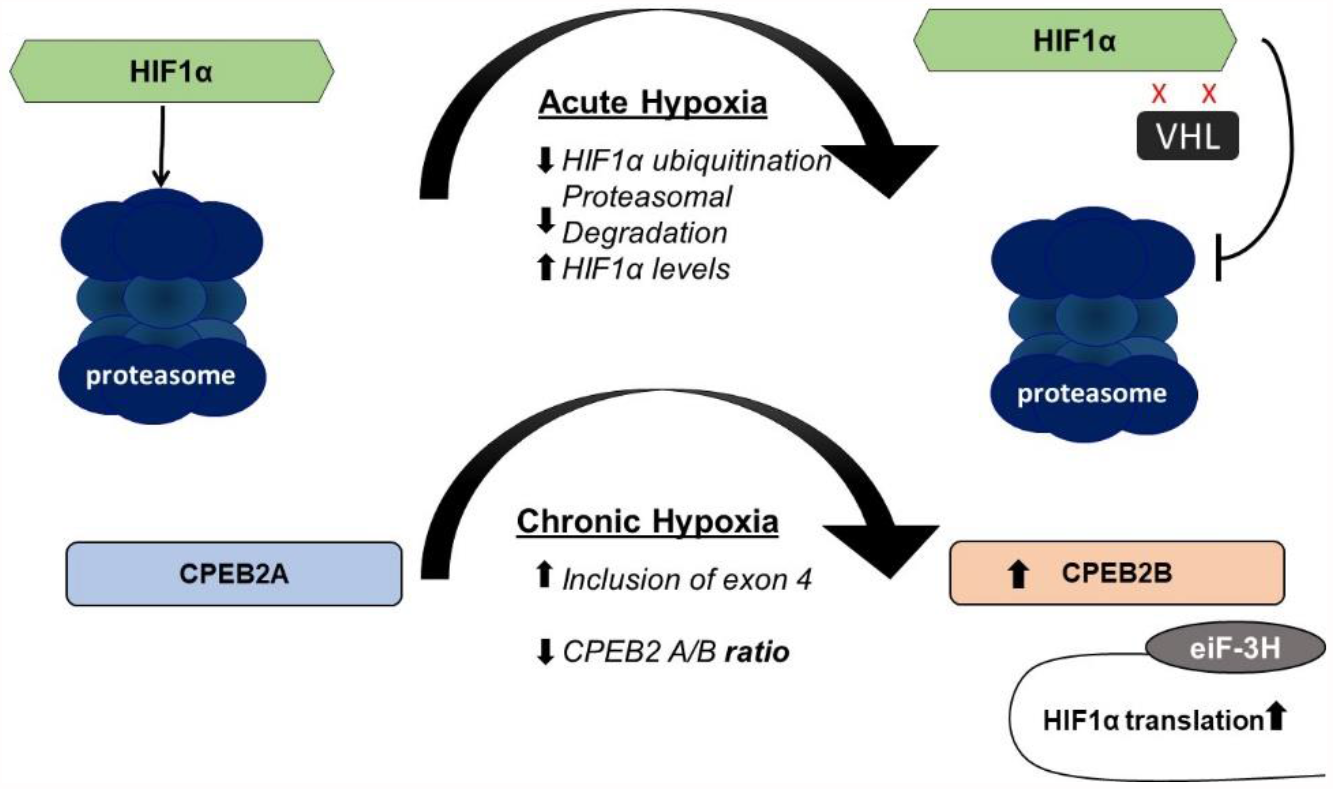
Schematic representation of CPEB2B-mediated HIF1α translation during chronic hypoxia.

We have demonstrated in Figures 1-3 that HIF1α translation is controlled at least partially via CPEB2 alternative splicing. Thus, we wished to identify additional components of this pathway. Hutt et al (**21**) have demonstrated that HIF1α translation may be regulated by initiation factor eIF3H. We, therefore, wanted to determine if CPEB2B may be acting via this mechanism. We demonstrate in **Fig. 5E-F** that not only does inhibition of eIF3H via RNAi decrease CPEB2B-mediated HIF1α expression, but that this mechanism is likely due to translation as demonstrated in **Fig. 5F, S4**.

## DISCUSSION

Structural changes to the pulmonary vasculature as a result of chronic hypoxia can lead to elevated and irreversible increases in arterial pressure and downstream pathologies. Despite current treatment methods, pulmonary hypertension remains a prevalent condition characterized by thick arterial walls, high blood pressure, and shortened survival. By targeting the genes involved in the hypoxic response, the integrity of the arterial walls might be preserved, and disease progression alleviated. Consequently, understanding and characterizing the mechanisms behind this response could benefit the development of future therapeutics involved in hypoxia-induced pathologies.

It is well documented that CPEB family members bind to specific mRNA targets, thereby preventing translation (**19**). We find that the CPEB2B isoform has the opposite effect for some targets, inducing the translation of HIF1α and TWIST1 (**15**,**16**). Hence, the inclusion of the small 30 amino acid exon in the CPEB2B sequence dramatically changes the function of this protein (**15**,**16**). In both these and previous studies, we find that, in contrast to the function of the A isoform, expression of HIF1α during the hypoxic response is sustained at least in part by translation via the CPEB2B isoform. Previous findings from our laboratory indicate that the role of CPEB2B in cancer metastasis is key to the advancement of disease in triple negative breast cancer and that this increase in hypoxia-linked factors is due to translation (**14-16**). We hypothesize that the presence of the CPEB2B isoform is critical for nascent HIF1α protein production at later stage timepoints during oxygen deprivation and this is further supported by our data demonstrating that VEGF expression may be regulated via CPEB2 A/S. Our hypothesis to explain this phenomenon is that there is a transitional stage between acute and chronic phases of hypoxia and that shift leads to increased HIF1α translation and decreased degradation. One interesting finding is that HIF1α protein expression is increased basally after cycloheximide treatment. Hence, we hypothesize that the cycloheximide itself induces a small amount of stress. Indeed, it has been demonstrated that cycloheximide can deplete ubiquitin in cell systems (**36**). It is therefore plausible that this depletion affects HIF1α levels basally.

With respect to our HUVEC spheroid assay, both sprout length and number were dramatically decreased after hypoxia when CPEB2B is inhibited. However, we did note that, when treated with CPEB2B ASO, sprout *diameter* seemed to be increased. Although we cannot say for certain why the CEB2B ASO increased sprout diameter, we hypothesize that a physical explanation is possible. It may be that the large number of sprouts induced by hypoxia in our control spheroids made it such that there was no physical room for larger diameter sprouts. Furthermore, the spheroids derived from HUVECs treated with CPEB2B ASO did not grow in culture at the same rate as control and were, therefore, smaller as shown in Figure 2. This observation is in general agreement with our findings in cancer models that CPEB2B over, in contrast to the A isoform, greatly increases tumor size and metastasis (**15-16**). Furthermore, we find that CPEB2B ectopic expression leads to small changes in CSL, but no significant changes in sprout number per spheroid. These data lead us to postulate that the increase in endogenous CPEB2B is likely sufficient to enhance VEGF signaling in this model. Interestingly, we find minimal changes in CSL and sprout number when we ectopically express CPEB2A (exon excluded), indicating that CPEB2B, and not CPEB2A function may be required in this case.

Regarding the mechanism of function of CPEB2B, there are multiple possibilities for why it is different than that of CPEB2A. We find that the proteins seem to bind differentially to UUUUUAU consensus sequences based on the surrounding sequence (based on our findings in **Fig.5**). This finding is suggestive of a possible shift in the RNA binding domain to bind more readily to other sequences surrounding the consensus CPE site. Analysis of the binding domain of CPEBs by Richter and colleagues suggests that both RNA recognition motifs as well, as the N-terminal zinc finger domain are required for binding (**22**). Furthermore, RNA footprinting studies by this group have elegantly demonstrated that the secondary structure of RNA is important for the binding of CPEB family members (**23**). Yet others have demonstrated a “fly-trap” mechanism of RNA binding whereby the two RRM domains enclose the RNA sequence (**24**). other sequences that are important for CPE recognition include the polyadenylation sequences. However, we do not observe a polyadenylation consensus sequence in proximity to the exon 10 site. Hence, we hypothesize that changes in the N-terminal domain causes changes in the secondary structure of the RNA which is recognized by CPEB2. The global sequences chosen for binding by CPEB2 isoforms is an active topic of research in our laboratory. Regarding exon 4 of CPEB2, the structure of RNA-bound CPEB2B RRMs is currently unknown.

However, IUPRED and Anchor analysis of the sequence of CPEB2A and CPEB2B indicates that the included exon is likely a protein-protein interaction domain and is a slightly intrinsically disordered to slightly ordered region (**Fig.S3**). Hence, it is possible that CPEB2B targets bound mRNA species to the translational machinery via protein-protein interactions. Chen et al. (**25**) have identified an eEF2 interacting domain N-terminal to the included exon for CPEB2B, which is responsible for CPEB2A-mediated inhibition of expression of HIF1α. It is thus possible that CPEB2B binds to other factors which then mediate eIF3H-dependent translation of mRNAs and we will pursue this possibility in future studies.

In conclusion, we have demonstrated that a CPEB2B-mediated translational mechanism may also play an important role during the chronic phase of the hypoxic response pathway. Our data indicate that CPEB2B may cooperate with eIF3H to induce translation of the HIF1α mRNA during the chronic phase of the cellular hypoxic response. However, it is not known precisely how this happens. Thus, future directions for this research include animal studies and studies to determine the factors that direct sequence and secondary structure choice when binding to RNA. HIF1 signaling plays a major role in the pulmonary response to chronic hypoxia. Therefore, patients with adverse pulmonary outcomes secondary to the chronic hypoxic state may benefit from these findings after future studies have been completed.

## MATERIALS AND METHODS

### Cell culture

All cell lines (HUVEC (CRL-1730), hPAEC (PCS-100-022), HEK293 (CRL-1573)) were purchased from ATCC (American Type Culture Collection) and cultured according to manufactures instructions. Both endothelial cell lines (HUVEC, hPAEC) were grown in EBM-2 media with EGM-2 bullet kit (purchased from Lonza). No additional serum was used for cell culture. HEK393 cells were grown in DMEM (10% FBS, 5% P/S). All experiments were performed on HUVEC’s and hPAEC’s ranging from passage 1 to passage 6. When needed, cells were split with Trypsin-EDTA (0.25%).

### Antibodies and reagents

Two different HIF1α (105SS and 610958) antibodies were used from Novus Biotechnologies and BD Transduction Laboratories. Anti-VEGF (MA513182) was purchased from ThermoFisher. Cycloheximide (C7698) was purchased from Sigma-Aldrich. Click-it protein reaction buffer kit was purchased from Thermofisher. Biotin for click it reaction was purchased from Sigma. All primers and RNA sequences used in this study were purchased from Integrated DNA Technologies (IDT). The HA-HIF1alpha-pcDNA3 plasmid was a kind gift from Dr. William Kaelin (Addgene plasmid #18949).

### Hypoxia exposure

A Modular Incubator Chamber (MIC-1) was purchased from Billups-Rothenberg Inc. Custom hypoxia mix (2% oxygen, 5% carbon dioxide, balance nitrogen, size 200 certified standard-spec cga 580) was purchased from Airgas USA LLC. Cells plated on either 10cm-dishes or 6-well plates (Corning, Sigma Aldrich) were placed in the chamber with humidified air. Hypoxic air was flushed into the tank for 3-5 minutes, sealed, and added to a 37°C incubator for 10-15 minutes before re-administering hypoxic air for another 3-5 minutes. Cells were left in the chamber and incubated at 37°C for indicated times.

### Antisense oligonucleotide and plasmid transfection

ASOs and plasmids were both transfected using Lipofectamine 2000 reagent (11668019) purchased from Thermo-Fisher. Each transfection master mix was incubated in Opti-MEM for 20 minutes prior to adding to serum-starved cells. Cells were then incubated [overnight for plasmid transfection and co-transfection, 30 hours for ASO transfection] before replacing media with complete medium.

### Quantitative “Splicing” RT-PCR

cDNA was synthesized using the Superscript III kit (Life Technologies) according to the manufacturers’ instructions. cDNA libraries were subjected to traditional PCR as described previously (**16, 26, 27**) using primers located on either side of exon 4 of the CPEB2 gene as described previously (16). Ratios were then calculated using densitometry/ImageJ as described by us previously (**15-17**).

### Immunoblotting

Total protein (15–30 μg) was electrophoretically separated on 10% polyacrylamide gels. Samples were transferred electrophoretically to PVDF membranes, then probed with the appropriate antibody as described previously (**16, 26, 27**).

### Immunoprecipitation

Indicated proteins were immunoprecipitated as described previously using anti-FLAG antibodies (**15-17**).

### Streptavidin-biotin affinity pull-down assay

Protocol was adapted from Chen et al, 2012 (**25**). Biotinylated RNA HIF1α 3’ UTR sequences surrounding the CPE site were purchased from IDT Cells transfected with CPEB2A-flag tagged or CPEB2B-flag tagged were lysed via freeze-thaw in binding buffer. FLAG-tagged CPEB2 isoforms were immunoprecipitated as described previously (**15**,**16**) then incubated with biotinylated RNA sequences in binding buffer (100mM KCL, 10mM HEPES, 0.1mM CaCl_2_, 1mM MgCl_2_, 5% glycerol, 100uM ZnCl_2,_ 0.1 mg/mL BSA, and 1x protease phosphate cocktail and 1x RNAse out, plus 5mM DTT and tRNA). Samples were incubated at room temperature for 30 mins and allowed to rotate overnight with streptavidin beads (washed in binding buffer). Beads were then washed in 1x binding buffer three times and proteins were eluted Laemmli buffer.

### Electrophoretic mobility shift assay

FITC-conjugated RNA sequences were incubated with immunoprecipitated CPEB2A-FLAG protein in binding buffer (**16**) then electrophoresed on a 5% acrylamide gel in 0.5 x TBE. Fluorescent bands were visualized.

### siRNA Transfection

Custom and validated siRNA targeted towards CPEB2B utilized in this study were introduced into each cell sample as described previously (**16**).

### Nascent protein labeling

Protocol was adapted from DeLigio et al, 2017 **(15)**. Cells were incubated in RPMI medium with 200 μM L-azidohomoalanine for 5 hours. Cells were then harvested as previously described and lysed in lysis buffer (1% NP-40 in 50 mM Tris-HCl, pH 8.0, 1 x protease and phosphatase inhibitors, added fresh). Cells were then sonicated and centrifuged to clear the lysate. Nascent proteins were labeled with biotin using the Click-it protein reaction buffer kit (Thermo Fisher) according to the manufacturer’s instructions. Nascent proteins were precipitated using streptavidin-coated magnetic beads. Following 4 washes in PBS 7.4 pH +/- 1% NP40, samples were subjected to PAGE-immunoblot.

### HUVEC spheroid generation

HUVEC spheroids of defined cell number were generated as previously described **(28, 29)**. In brief, HUVECS below passage 7 were grown to 75-90% confluency and trypsinized. Cells were re-suspended in EGM-2 Lonza medium containing 20% methylcellulose and cultured as hanging drops (500 cells / 25µL) onto the lid of a petri dish and incubated at 37 C overnight to form spheroids.

### Transfection for HUVEC spheroids

HUVEC’s were grown to 75-80% confluency in 75cm^2^ flasks before transfection with ASO-B, c-ASO, CPEB2A or CPEB2B plasmid. Lipofectamine 2000 was used according to the manufacturer’s instructions to introduce the appropriate ASO or plasmid to the monolayer of cells. For each transfection, cells were incubated with the lipofectamine in a serum-free OPTI-MEM media overnight before being trypsinized with Trypsin-EDTA (.25%). Each transfected group of cells were then generated into spheroids as described above.

### *In vitro* sprouting assay

Approximately 100 HUVEC spheroids were collected by gently washing with 10x PBS and centrifuging at 500g x 5 min. On ice, rat-tail collagen was mixed with DMEM (8:1) and NaOH (enough to change the solution from yellow to faint pink). Spheroids were resuspended in EGM-2 Lonza media supplemented with 30% FBS, and 0.25% methylcellulose. Immediately following the addition of the collagen:DMEM mixture to the spheroids, 500uL of each spheroid mixture was rapidly plated into a 24-well plate and allowed to polymerize at 37C for 30 minutes. After polymerization, cells were exposed to hypoxia as described above. Sprout number per spheroid was counted, as well as cumulative sprout length using the measurement tool in FIJI/ImageJ.

