## Supplement for "Chronic hypoxia regulates Cytoplasmic Polyadenylation Element Binding Protein 2 alternative splicing to promote HIF1a translation"

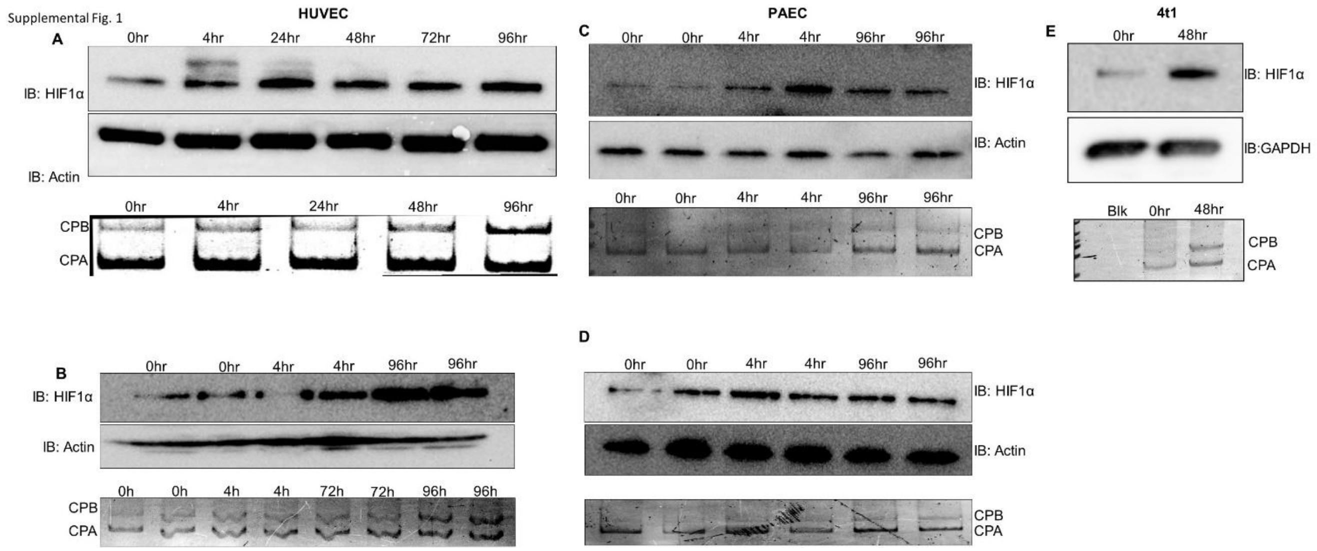


**Figure S1: CPEB2 A/S is dysregulated during chronic hypoxia.** A-D: Immunoblots and competitive PCR gels represent biological replicates of blots found in Fig.1. C: Immunoblots and competitive PCR gels representing the same phenomenon in a murine cell line.

**Supplemental Fig.2**

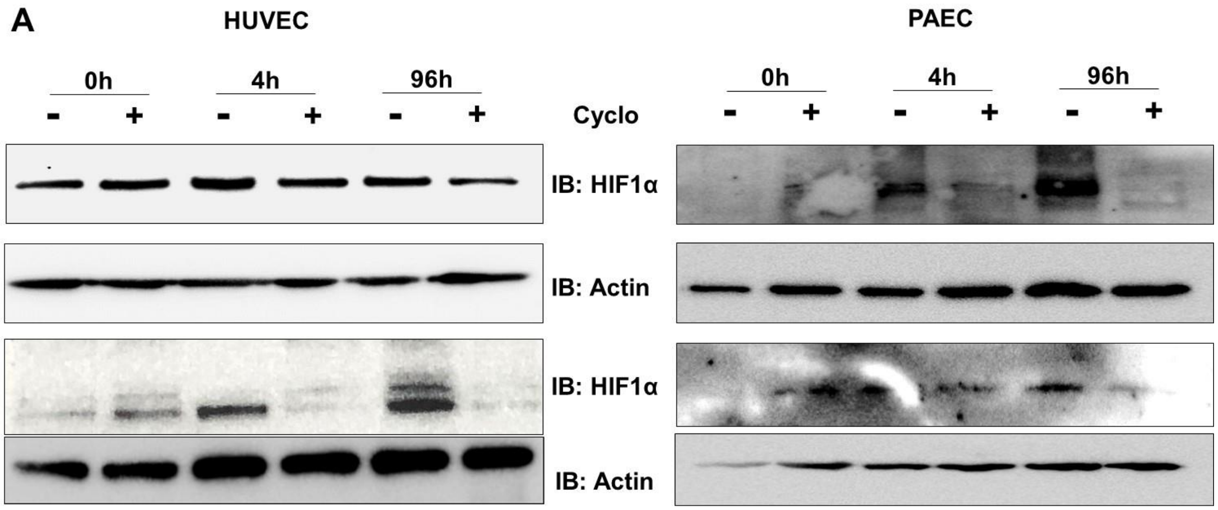


**Figure S2: Translation of HIF1α during chronic hypoxia.** A: Immunoblots represent biological replicates of blots found in Fig.3.

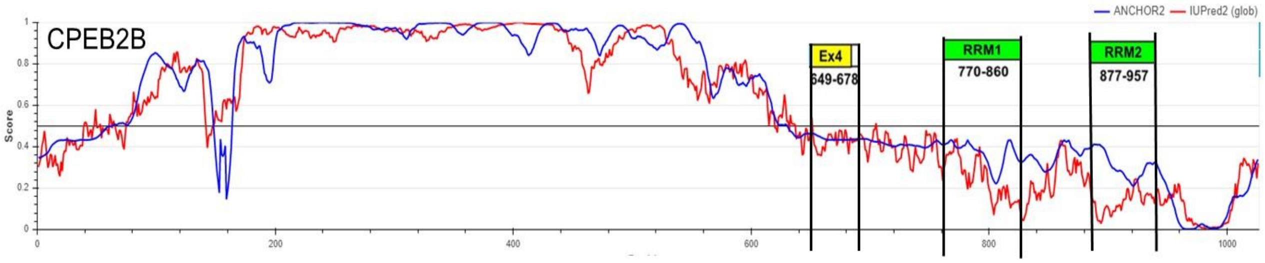


**Figure S3: CPEB2 exon 4 is in a slightly ordered region predicted to be a protein-protein interaction domain.** IUPRED (red) and Anchor (blue) scores of CPEB2B protein sequence.

**Supplemental Figure 4**

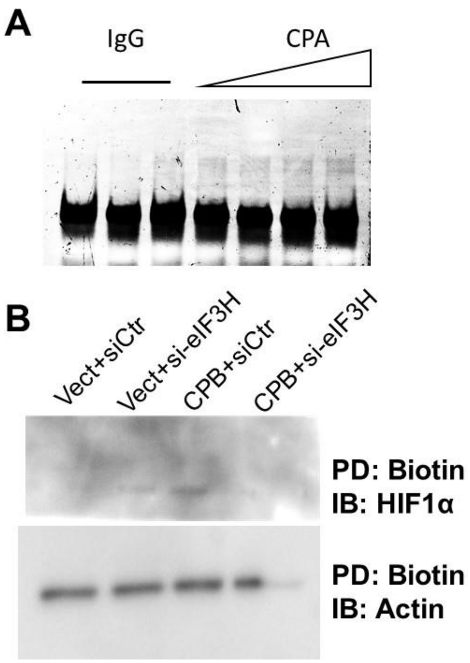


**Figure S4: CPEB2A binds to the HIF1α 3'UTR CPE site directly and subsequent translation of HIF1α is eIF3H-dependent.** A: EMSA of CPEB2A in increasing concentrations using the HIF1α 3'UTR CPE sequence as a probe. B: CLICK nascent protein assay under the indicated conditions.
